# An MLL-Independent Function of Menin Promotes Resistance to MAPK-Targeted Therapy

**DOI:** 10.64898/2026.08.24.746076

**Authors:** Akash Srivaths, Aleen Al Halawani, Tirta M Djajawi, Anne Huber, Chloe Gerak, Laura Jenkins, Rebekah Crake, Kristen A Needham, Biswadeep Sen, Isela Sarahi Rivera, Nooshin Khoshdoozmasouleh, Lisa A Mielke, Liam Neil, Bhupinder Pal, John M Mariadason, Conor J Kearney, Stephin J Vervoort

**Affiliations:** Olivia Newton-John Cancer Research Institute, Heidelberg, VIC, 3084, Australia; School of Cancer Medicine, La Trobe University, Melbourne, VIC, 3086, Australia; The Walter and Eliza Hall Institute of Medical Research, Parkville, VIC, 3052, Australia; La Trobe Institute for Molecular Science, La Trobe University, Bundoora, VIC, 3086 Australia

## Abstract

BRAF mutant colorectal cancer (CRC) remains difficult to treat despite the clinical use of combined BRAF and EGFR inhibition, highlighting a need to define tumour-intrinsic mechanisms that limit therapeutic response. Here, using genome-wide CRISPR-Cas9 screening in BRAF-mutant CRC cells, we identify MEN1, encoding the chromatin-associated protein Menin, as a selective determinant of sensitivity to combined encorafenib and cetuximab (EC). MEN1 loss markedly enhanced EC-mediated inhibition of cell proliferation and ERK activity while having comparatively little effect in untreated cells, and re-expression of Menin restored resistance. Transcriptomic and chromatin profiling revealed that Menin supports the transcriptional response associated with MAPK signalling. Menin occupied promoters of MAPK/BRAF-responsive genes and EC treatment caused widespread displacement of Menin from chromatin. Phosphoproteomic analysis demonstrated extensive remodelling of MAPK signalling following EC treatment, whereas proximity proteomics showed that the Menin-associated protein complexes remained largely intact despite loss of Menin chromatin occupancy. Importantly, MLL1 loss did not reproduce the sensitising effect of MEN1 deletion, and pharmacological Menin inhibition with revumenib failed to phenocopy either genetic MEN1 loss or acute Menin degradation, indicating that this phenotype is independent of Menin-MLL activity. Together, these findings identify a previously unrecognised, MLL-independent role for Menin in buffering the response of BRAF-mutant CRC cells to MAPK pathway inhibition and suggest targeting Menin, rather than disruption of its interaction with MLL, may provide a strategy for enhancing the response to BRAF-targeted therapy for CRC.

## Introduction

BRAF V600E mutations occur in approximately 8-12% of metastatic colorectal cancers and define a subtype with markedly poor prognosis^1^. These tumours respond poorly to BRAF inhibition alone because of rapid pathway reactivation through EGFR^2,3^. Combined inhibition of BRAF and EGFR is therefore required^4,5^. Based on the BEACON CRC trial, encorafenib plus cetuximab (EC) is approved for previously treated BRAF V600E metastatic colorectal cancer and is the only chemotherapy-independent targeted regimen available for these patients^6,7^. However, the confirmed objective response rate was only 19.5%, with a median overall survival of 9.3 months, compared with 5.9 months for chemotherapyInherent^8^ and acquired resistance underpins the limited efficacy of this treatment.

CRISPR screening has emerged as a powerful tool for dissecting genes that when deleted, provide sensitization or resistance to diverse cellular stimuli^9^. We performed a genome-wide CRISPR knockout screen in BRAF V600E RKO colorectal cancer cells treated with EC. Parallel screens were performed with an ERK inhibitor or 5-fluorouracil to distinguish vulnerabilities associated with MAPK pathway inhibition from more general effects on drug sensitivity. RKO cells were selected as prior studies have shown them to harbor *de novo* resistance to BRAF inhibition^10^. Indeed, Menin was one of the strongest sensitiser genes identified. Menin, encoded by MEN1, is a nuclear scaffold with no catalytic activity of its own^11^. Its function varies considerably between cellular contexts. Germline loss-of-function mutations in MEN1 cause multiple endocrine neoplasia type 1, and in endocrine tissues Menin acts as a tumour suppressor^12^. It interacts with MLL to promote expression of the cyclin-dependent kinase inhibitors p18INK4c and p27Kip1^13^ and can directly repress the AP-1 factor JUND^14^. In MLL-rearranged leukaemia, Menin instead has an oncogenic role, anchoring MLL fusion proteins and LEDGF to chromatin at genes including HOXA9 and MEIS1^15,16^. The same Menin binding pocket can interact with MLL and JUND despite their different transcriptional consequences^17^. Menin has also been linked to colorectal cancer. Menin inhibition has been reported to repress SKP2 and increase sensitivity to EGFR inhibitors^18^, while Menin-dependent repression of glycolysis has also been implicated in resistance to EGFR inhibition^19^.

Current Menin inhibitors were developed to disrupt its interaction with MLL. Revumenib, ziftomenib, bleximenib, icovamenib and enzomenib bind the Menin-MLL pocket and disrupt the association of Menin with MLL fusion proteins. These compounds have shown activity in MLL-rearranged and NPM1-mutant acute leukaemia^20,21^ and revumenib is now approved for acute leukaemia^22^. This approach works because the Menin-MLL interaction is required in these cancers. Whether inhibiting the same interaction is sufficient to target Menin dependencies in solid tumours is much less clear.

Here, we show that loss or targeted degradation of Menin sensitises BRAF V600E colorectal cancer cells to EC. In contrast, deletion of MLL or treatment with revumenib does not reproduce this effect. EC substantially reduces Menin occupancy on chromatin without disrupting the broader Menin protein complex, while revumenib affects only a small fraction of Menin-associated proteins. The dependency identified by the CRISPR screen therefore does not appear to reflect the canonical Menin-MLL interaction. Instead, our findings point to a Menin function required during MAPK pathway inhibition that is lost when Menin is removed but retained when its MLL-binding pocket is inhibited. This distinction is important for Menin and potentially other non-catalytic scaffold proteins, where genetic loss of the protein and pharmacological inhibition of a single interaction may produce very different effects.

## Results

### MEN1 is not essential in CRC but is the top synthetic-lethal sensitiser to EC

To identify genes whose loss potentiates sensitivity to MAPK pathway inhibition, we performed a genome-wide CRISPR knockout screen in RKO (BRAF V600E) cells (**Figure 1a**). Cas9-expressing cells were transduced with a genome-wide sgRNA library, selected, and split across three treatment arms alongside an untreated time-matched control: encorafenib plus cetuximab (EC), the ERK inhibitor AZD0364 (ERKi), and 5-fluorouracil (5-FU). EC represents the current clinical standard of care for BRAF V600E CRC; ERKi tests whether hits are specific to inhibition of the MAPK axis more broadly; and 5-FU serves as a cytotoxic specificity control.

**Figure 1.**
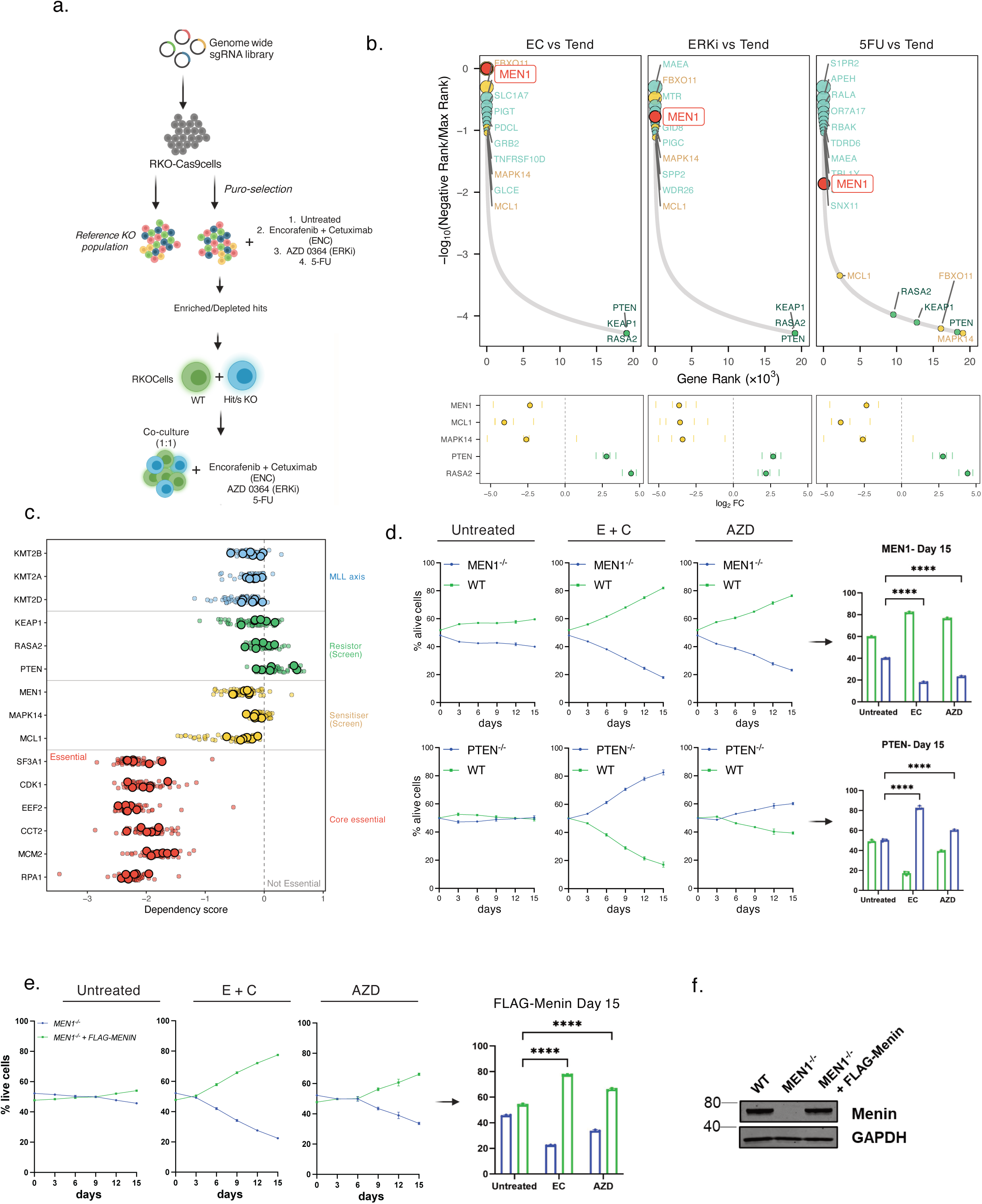
Genome-wide CRISPR screening identifies MEN1 as a sensitiser to MAPK pathway inhibition in BRAF-mutant colorectal cancer. (a) Schematic of the genome-wide CRISPR-Cas9 knockout screen performed in RKO BRAF V600E colorectal cancer cells. Cas9-expressing cells were transduced with the Brunello genome-wide sgRNA library and divided into untreated, encorafenib plus cetuximab (EC), AZD0364 (ERKi) and 5-fluorouracil (5-FU) treatment arms. sgRNA abundance was determined at the experimental endpoint and compared with the time-matched untreated population. (b) CRISPR screen results for EC, ERKi and 5-FU treatment. Upper panels show gene-level depletion ranked across each screen; genes are ordered by rank and the y-axis is -log10 of rank position. Lower panels show sgRNA-level log2FC for selected sensitiser and resistance genes; ticks represent individual sgRNAs and filled points the per-gene median. MEN1, MAPK14 and MCL1 were depleted following MAPK pathway inhibition, while PTEN, RASA2 and KEAP1 were enriched. (c) DepMap CRISPR Chronos dependency scores for MEN1, KMT2 family members, selected screen hits and core-essential genes across colorectal cancer cell lines. Larger points indicate BRAF V600E cell lines (RKO, H29, COLO205, LS411N, MDST8, HT115). (d) Competition assays comparing wild-type and MEN1-knockout or PTEN-knockout RKO cells under untreated, EC (500 nM and 20 µg/mL respectively)- and ERKi (250 nM)-treated conditions. The proportion of viable cells of each genotype was followed over time. Representative images from one of the three independent experiments (n=3) are shown. Data presented as mean ± SD. Statistical significance was determined using two-way ANOVA. **** p < 0.0001 (e) Competition assay following re-expression of FLAG-MENIN in MEN1-knockout RKO cells under EC (500 nM and 20 µg/mL respectively) treatment. Representative images from one of the three independent experiments (n=3) are shown. Data presented as mean ± SD. Statistical significance was determined using two-way ANOVA. **** p < 0.0001 (f) Immunoblot confirming restoration of Menin expression following FLAG-MENIN re-expression.

*MEN1* was the most depleted gene under EC, ranked first of 19,113 (log2FC -2.36, FDR 0.077), and ranked sixth under ERKi (log2FC -3.64, FDR 8.3 x 10^4^). Under 5-FU it ranked 73rd and did not reach significance (log2FC -1.76, FDR 0.70), indicating specificity for MAPK pathway inhibition rather than general cytotoxic stress (**Figure 1b, upper panels**). Only seven genes were called as sensitisers in the EC arm at FDR < 0.25, against 118 resistors, and *MEN1* was the most depleted. Depletion was consistent across independent sgRNA for *MEN1*, *MCL1* and *MAPK14*, while guides targeting *PTEN* and *RASA2* were uniformly enriched, validating these as resistors (**Figure 1b, lower panels**).

The identity of the other hits is consistent with the drug’s mechanism. *MAPK14* (encoding p38-alpha) provides an alternative stress-activated MAPK survival signal when the canonical ERK cascade is blocked, and its loss removes this backup^23^. *MCL1* encodes an anti-apoptotic BCL-2 family member that sequesters BAX and BAK^24^. Its identification as a sensitiser suggests that EC-treated cells already sit near an apoptotic threshold that MCL1 buffers. The top resistors (*PTEN*, *RASA2* and *KEAP1*) define the escape routes available under EC. *PTEN* loss hyperactivates PI3K/AKT, a critical survival axis when MAPK signalling is inhibited^25^. *RASA2* encodes a RAS GTPase-activating protein whose loss sustains RAS in its active state^26^, potentially re-wiring signalling around the BRAF and EGFR block. *KEAP1* loss stabilises NRF2 and raises oxidative stress tolerance^27^. The screen therefore reports coherent pathway biology rather than generalised drug tolerance, and places *MEN1* at the head of it.

To determine general dependency or a context-specific vulnerability, we analysed DepMap CRISPR dependency scores across CRC cell lines^28^. *MEN1* and the MLL family members it scaffolds including *KMT2A (MLL1)*, *KMT2B* and *KMT2D* all clustered near zero, comparable to the validated screen sensitisers (*MCL1*, *MAPK14*) and resistors (*PTEN*, *RASA2*, *KEAP1*), and far from core essential genes (*RPA1*, *MCM2*, *CCT2*, *EEF2*, *CDK1*, *SF3A1*) (**Figure 1c**). This held across BRAF V600E lines including RKO, establishing that Menin loss is tolerated in unperturbed CRC cells. *MEN1* is therefore not essential, but a synthetic-lethal partner.

We next performed competition assays to confirm the screen results. *MEN1*-knockout RKO cells were selectively outcompeted under EC, falling from 80% to 20% of the live population by day 15, but remained at parity with wild-type cells in untreated co-culture, while *PTEN*-knockout cells were enriched under EC, consistent with PI3K-mediated resistance (**Figure 1d**). The sensitisers *MAPK14* and *MCL1* (**Supplementary figure 1a)**, and the resistors *RASA2* and *KEAP1,* to a lesser extent, (**Supplementary figure 1b**), also showed consistent phenotypes across the EC and ERKi arms. Reintroduction of FLAG-MENIN into *MEN1*-knockout cells rescued the EC sensitisation phenotype, with FLAG-MENIN cells rising to 80% of the population by day 15 compared with 20% for *MEN1*-knockout cells (**Figure 1e**), and restored Menin protein was confirmed by immunoblot (**Figure 1f**). The sensitisation to EC can therefore be attributed to loss of Menin.

### EC suppresses MAPK signalling and Menin loss potentiates this effect

Having established the genetic interaction, we asked what each perturbation does transcriptionally. To associate transcriptional changes with upstream signalling we used PROGENy, which infers pathway activity from the coordinated behaviour of experimentally derived target gene sets rather than from the expression of pathway components themselves ^29^. The distinction matters here because kinase cascades are regulated post-translationally, and their own transcripts move little under inhibition. EC treatment alone produced a response dominated by a single axis. As expected, MAPK activity was suppressed dramatically, compared to any other pathway, with the related EGFR pathway second (**Figure 2a left panel**). These data confirm the EC drug combination disrupts the action of the intended signalling cascade.

**Figure 2.**
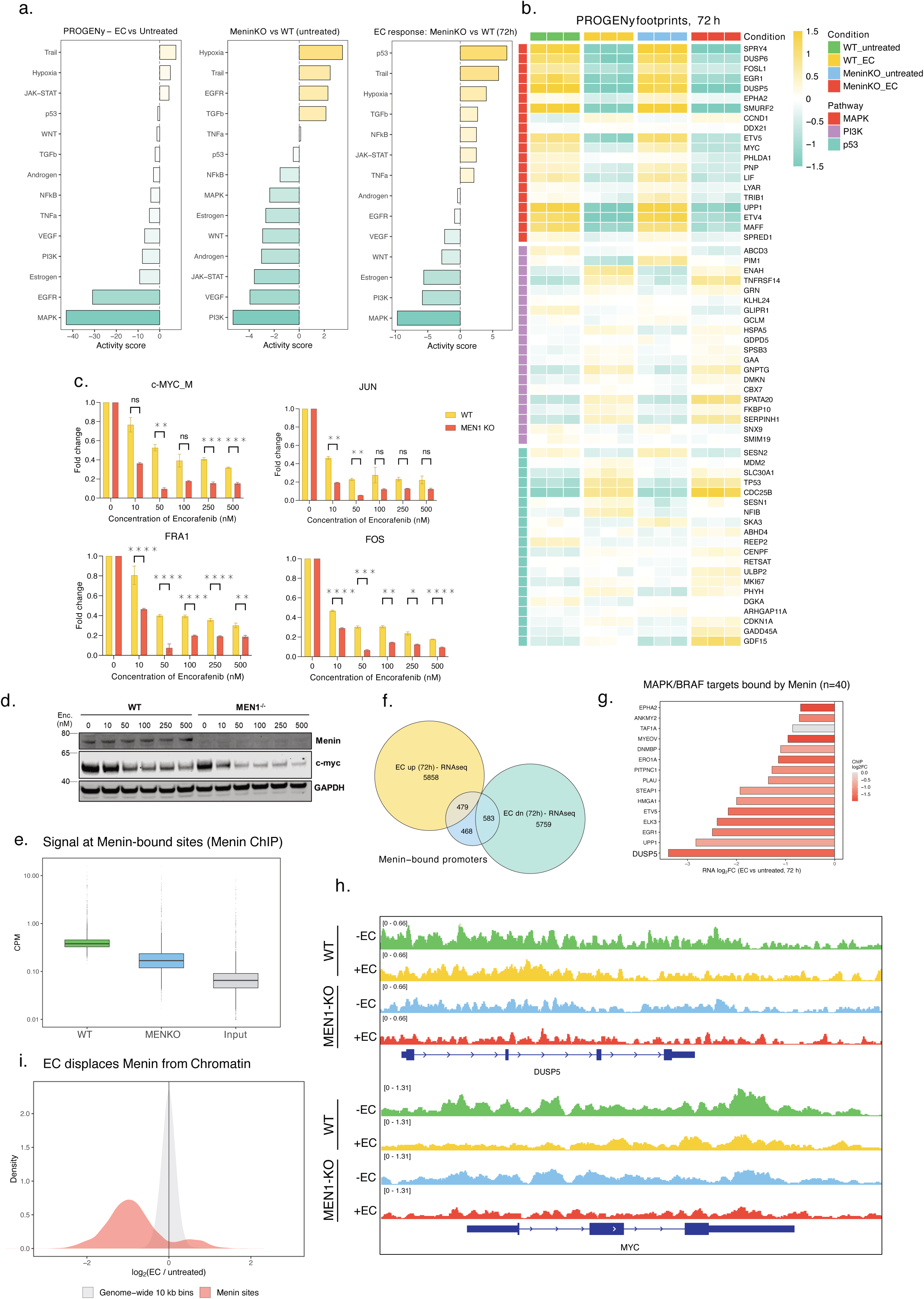
Menin regulates the transcriptional response to MAPK pathway inhibition and is displaced from chromatin following EC treatment. (a) PROGENy pathway activity scores derived from RNA-seq data for EC-treated versus untreated wild-type cells, untreated MEN1-knockout versus wild-type cells, and EC-treated MEN1-knockout versus EC-treated wild-type cells at 72h. (b) Heatmap showing expression of genes contributing to MAPK, PI3K and p53 pathway activity across wild-type and MEN1-knockout RKO cells with or without EC treatment at 72h. Expression values are shown as mean-centred log2 CPM. (c) MYC, JUN, FRA1 and FOS transcript abundance measured across increasing concentrations of encorafenib in wild-type and MEN1-knockout cells treated with cetuximab (20 µg/mL). Representative image from one of the two independent experiments (n=2) is shown. Data presented as mean ± SD. Statistical significance was determined using two-way ANOVA. ns = not significant, ** p < 0.01, *** p < 0.001 (d) Immunoblot analysis of c-MYC protein in wild-type and MEN1-knockout cells following increasing concentrations of encorafenib in the presence of cetuximab. GAPDH, loading control. (e) Menin ChIP-seq signal at Menin-dependent binding sites identified by comparison of wild-type and MEN1-knockout RKO cells. Wild-type-specific peaks were retained as Menin-dependent sites (n = 3,377). Signal is shown as CPM. (f) Overlap between genes with a Menin-bound promoter (n = 1,607) and genes significantly up- or downregulated following EC treatment at 72h in WT cells (p.adj < 0.05). Promoter binding was defined as a Menin-dependent site within 2 kb of the transcription start site. (g) MAPK/KRAS-responsive genes with Menin-bound promoters, showing changes in gene expression and Menin occupancy following EC treatment (6h). Bar, transcript log2FC; fill, menin ChIP log2FC at promoter. (h) Representative Menin ChIP-seq tracks at the DUSP5 and MYC loci in wild-type and MEN1-knockout cells with or without EC treatment. (i) Distribution of EC-induced changes in Menin ChIP-seq signal across Menin-dependent sites compared with genome-wide 10-kb bins. EC reduced Menin occupancy across Menin-dependent sites, with little change across the genome-wide background. Dashed line, median at Menin dependent sites. ChIP-seq was performed at n = 1 per condition.

Menin loss alone produced a different profile. PI3K was the most strongly suppressed axis, and while MAPK was also reduced it was neither the dominant effect nor of a magnitude comparable to that produced by EC alone (**Figure 2a middle panel)**. Loss of Menin is therefore not equivalent to a partial dose of MAPK inhibition.

The combination was not simply additive. In *MEN1*⁻/⁻ cells treated with EC for 72h, MAPK suppression exceeded that produced by either perturbation alone, and p53 emerged as the dominant induced axis, a pathway that moved negligibly under EC alone or under Menin loss alone (**Figure 2a right panel**). Menin loss thus potentiates the response to MAPK inhibition while adding a second arm that neither perturbation generates by itself.

Examining the genes carrying these pathway scores brought the effect down to individual targets (**Figure 2b**). *MYC*, the canonical transcriptional endpoint of the BRAF/MEK/ERK axis, was suppressed by EC in wild-type cells and suppressed further in *MEN1*⁻/⁻ cells treated identically, despite indistinguishable baseline expression between genotypes. The additional suppression was confirmed at transcript level by qPCR across an encorafenib dose range, where *MEN1*⁻/⁻ cells showed significantly lower *MYC JUN, FRA1 and FOS* at at the transcript level (**Figure 2c**), and MYC protein level by immunoblot, where c-Myc was progressively lost with increasing drug and further reduced in the absence of Menin (**Figure 2d**).

Among p53 pathway targets, *GADD45A* induction was substantially greater in *MEN1*⁻/⁻ cells, whereas *TP53* itself was strongly induced by EC in both genotypes and to a similar extent (**Figure 2b**). The inferred increase in p53 activity in Menin-deficient cells therefore reflects the behaviour of p53 target genes rather than of *TP53* transcription. PROGENy scores activity from the coordinated output of a weighted regulon, and a pathway can become more transcriptionally active without any change in the abundance of the transcription factor driving it. Menin loss increases what p53 does under EC, not how much p53 is produced.

### Menin occupies the promoters of EC-responsive genes and is displaced by treatment

The transcriptional data placed Menin loss in the same pathway space as EC without indicating where Menin acts. We therefore mapped Menin genome-wide by ChIPseq in wild-type and *MEN1*⁻/⁻ RKO cells.

Peaks called in *MEN1*⁻/⁻ cells cannot represent Menin, and we used them to define antibody background. Of 4,919 peaks stringently called in wild-type cells, 1,542 were also present in the knockout and were discarded; the remaining 3,377 wild-type-specific peaks were retained as Menin-dependent sites (**Figure 2e**). Signal at these sites was 2.3x higher in wild-type than in *MEN1*⁻/⁻ cells and 6.7x above input. Assigning sites to the nearest transcription start site identified 1,607 genes with a Menin-bound promoter (within 2 kb of the TSS).

Intersecting these genes with the transcriptional response to EC at 72h revealed substantial overlap in both directions: 583 Menin-bound promoter genes were significantly downregulated by EC and 479 were upregulated (**Figure 2f**). EC alters the expression of a large fraction of the transcriptome at this timepoint, and Menin binds preferentially at active promoters, which are themselves more likely to be drug-responsive.

Within the downregulated intersection, 40 genes were annotated targets of MAPK or KRAS signalling (**Figure 2g**). Restricting to canonical ERK target genes, five were Menin-bound with a 4x enrichment over the background rate of Menin binding among expressed genes. These were *DUSP5*, *EGR1*, *ELK3*, *ETV5* and *EPHA2*, and all five lost both Menin occupancy and transcript abundance under EC (**Figure 2g-h**). Menin was not enriched at the *MYC* promoter; six menin-dependent sites lie 63-116 kb downstream of the *MYC* TSS within the PVT1 locus; a long ncRNA which binds to and stabilises MYC, and lose Menin occupancy under EC (**Figure 2h**).

Displacement was not confined to these genes. Total menin protein was unchanged by EC treatment (**Figure 2d**), indicating that reduced occupancy reflects loss from chromatin rather than reduced protein abundance. Across all 3,377 Menin dependent sites, EC reduced Menin occupancy with a median log2FC of -0.94, compared with -0.01 across genome-wide 10 kb bins (**Figure 2i**). Displacement was greater at promoter-proximal than at distal sites (**Supplementary Figure 2a**) which suggests a graded relationship rather than a discrete separation into retained and displace classes. Furthermore, the magnitude of displacement at a given promoter did not predict the transcriptional response of the corresponding gene (Pearson r = -0.01, 95% CI −0.06 to 0.04, n = 1,530; **Supplementary Figure 2b**). Menin displacement and the EC transcriptional response therefore co-occur at these loci without being quantitatively coupled gene by gene.

### EC dismantles the signalling network around Menin without disrupting the Menin complex

The chromatin data established that EC removes Menin from DNA. We next asked whether it also alters the protein complex Menin assembles. We mapped the Menin protein neighbourhood by proximity labelling, expressing a doxycycline-inducible TurboID-Menin fusion and identifying biotinylated proteins after a 2h biotin pulse^30^ (**Figure 3a-b**). Proteins were retained only if significantly enriched over both a no-biotin and a no-doxycycline control (BH-adjusted p < 0.01, log2FC > 2), defining an interactome of 1,759 proteins. Of 1,025 *MEN1* interactors curated in BioGRID, 483 were detected in this proteome and 371 were retained in the stringent set, including the core MLL/COMPASS module (KMT2A, RBBP5, PSIP1, WDR5, HCFC1) and the AP-1 factors JUND and FOSL1 (**Figure 3c**).

**Figure 3.**
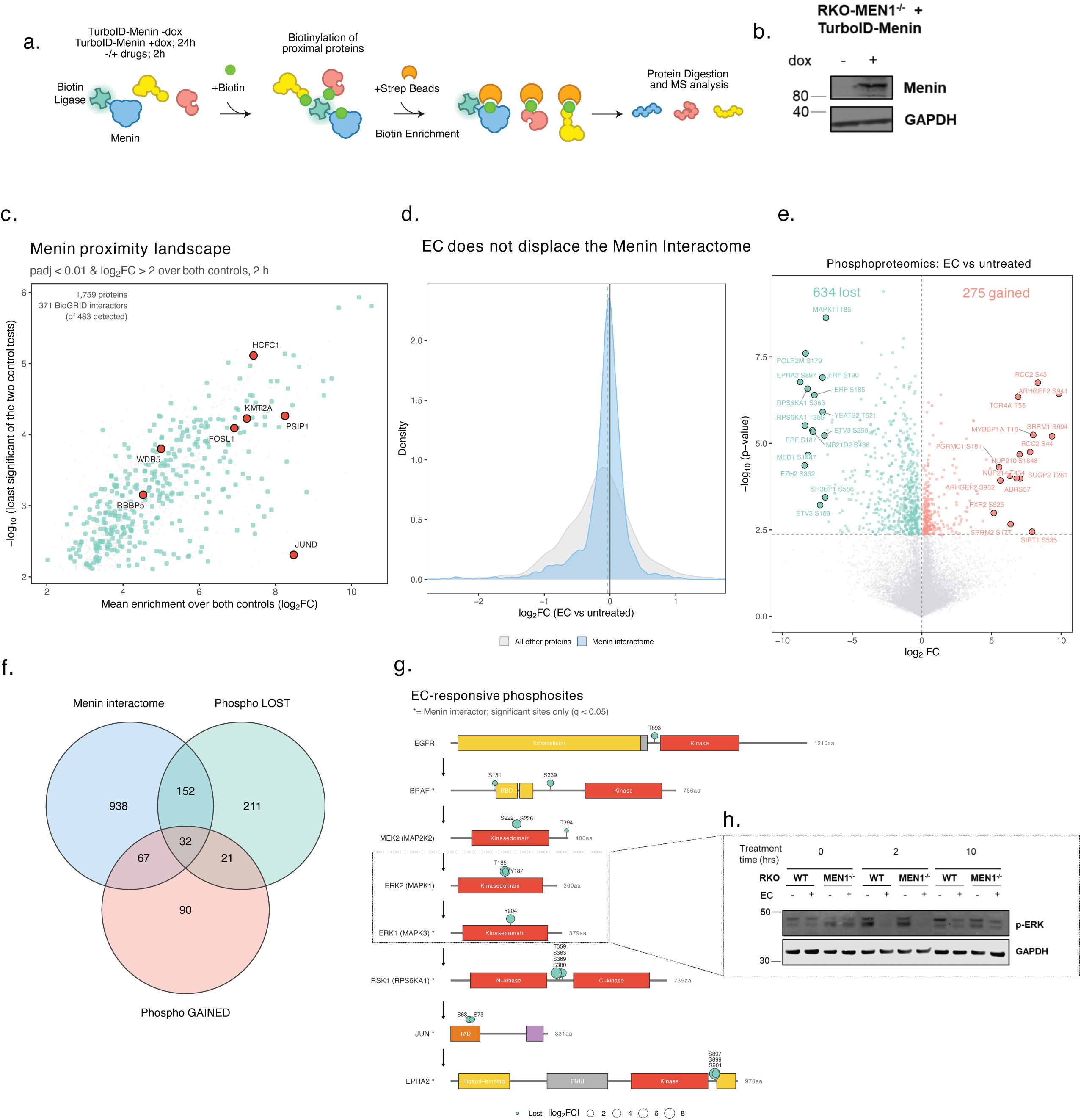
EC alters signalling associated with the Menin interactome without disrupting its overall composition. (a) Schematic of the inducible TurboID-Menin proximity-labelling approach used to define the Menin-associated proteome. Following doxycycline (1 µg/mL) induction, biotinylated proteins in proximity to TurboID-Menin were isolated by streptavidin enrichment and analysed by mass spectrometry. The construct was expressed in MEN1-KO background thus the fusion replaces rather than supplements endogenous menin. (b) Experimental conditions and controls used to identify proteins specifically enriched by TurboID-Menin proximity labelling. Proteins were retained in the Menin interactome when significantly enriched over both the no-biotin and no-doxycycline controls (BH-adjusted p < 0.01 and log2FC > 2 in each comparison), yielding 1,749 proteins. (c) Menin proximity landscape showing proteins enriched over both control conditions. Previously reported physical Menin interactors from BioGRID are indicated, including components of the MLL/COMPASS complex and AP-1 transcription factors. (d) Distribution of protein abundance changes following EC treatment for Menin interactors compared with all other quantified proteins. (e) Phosphoproteomic analysis of EC-treated versus untreated RKO cells. Phosphosites significantly increased or decreased following EC treatment are indicated (q < 0.05). (f) Overlap between the Menin interactome and proteins containing phosphosites significantly altered by EC treatment. Analysis is restricted to protein with at least one masured phosphosite. (g) EC-responsive phosphosites mapped across selected components of the EGFR-BRAF-MEK-ERK pathway and downstream effectors and ordered by position in the pathway. Domains are drawn to UniProt annotation and point size indicates the magnitude of the change; only sites with q < 0.05 are shown. Asterisks indicate proteins also identified in the Menin interactome. (h) Immunoblot analysis of ERK1/2 phosphorylation in wild-type and MEN1-knockout RKO cells over a time course of EC (500 nM and 20 µg/mL respectively) treatment (0, 2 and 10h). GAPDH, loading control.

EC treatment did not alter this network. The distribution of fold-changes at interactome proteins was indistinguishable from that across all quantified proteins **(Figure 3d)**. This revealed that despite Menin being removed from chromatin under EC, the protein complex around it remains largely intact.

We then asked whether the complex is instead modified. Phosphoproteomic profiling of EC-treated versus untreated cells identified 634 phosphosites significantly reduced and 275 increased (**Figure 3e**). Of the Menin interactors with a measured phosphosite, 184 lost and 99 gained phosphorylation (**Figure 3f**), indicating that EC alters the modification state of Menin’s neighbourhood extensively even as its composition remains unchanged. Overall, Kinase substrate enrichment analysis (KSEA) showed reduced inferred activity of MEK1, RSK1, RSK2, RSK3, TPL2, JAK2, BRAF and p38 targets, consistent with collapse of MAPK sganling cascade; and increased activity of CK2*α* as the only significantly activated kinase (Supplementary Figure 3c).

The sites lost mapped onto the drug-targeted cascade itself (**Figure 3g**). Phosphorylation was reduced at EGFR T693, BRAF S151 and S339, MEK2 S222 and S226, the ERK2 activation loop (MAPK1 T185, Y187) and the ERK1 activation loop (MAPK3 Y204), and continued into RSK1 (RPS6KA1 T359, S363, S369, S380), JUN (S63, S73) and EPHA2 (S897, S899, S901). Of these, BRAF, ERK1, RSK1, JUN and EPHA2 were themselves Menin interactors, whereas ERK2 was not.

Menin itself was not modified. Its single detected phosphosite, S487, was unchanged by EC (log2FC −0.09, q = 0.93). Nor was the core complex; of more than 100 phosphosites detected across MLL/COMPASS components, only four changed significantly SETD1A T1167 and S1169 (reduced) and KMT2B S1032 and S1035 (increased) and none of these were on Menin, KMT2A, RBBP5, ASH2L or PSIP1.

Immunoblotting for phospho-ERK (T202 and Y204) across a treatment time-course showed that ERK1 phosphorylation predominates in untreated wild-type cells and is markedly reduced at baseline in *MEN1*⁻/⁻ cells, with a corresponding increase in phospho-ERK2 (**Figure 3h**). *MAPK3* (ERK1) transcript was unchanged between genotypes (log2FC 0.08, adjusted p = 0.32), placing the difference downstream of transcription. At 2h, EC reduced phosphorylation of both isoforms in both genotypes; by 10h, total phospho-ERK remained lower in *MEN1*⁻/⁻ cells, with ERK2 more affected than ERK1.

### Sensitisation requires loss of Menin protein, not inhibition of its MLL-binding interface

Menin is targeted clinically by inhibitors of its interface with MLL. We asked whether that pharmacology reproduces the genetic effect.

Revumenib treatment displaced 71 of 1,674 quantified interactors from Menin (4.2%), leaving 1,603 (95.8%) in place (**Figure 4a).** The displaced fraction was coherent rather than random, comprising KMT2A with the COMPASS subunits RBBP5 and PSIP1, the AP-1 factors JUND and FOSL1, all three JADE subunits of the HBO1 acetyltransferase complex, and KMT5A. WDR5, HCFC1, JUNB and JUN remained associated with Menin despite KMT2A displacement.

**Figure 4.**
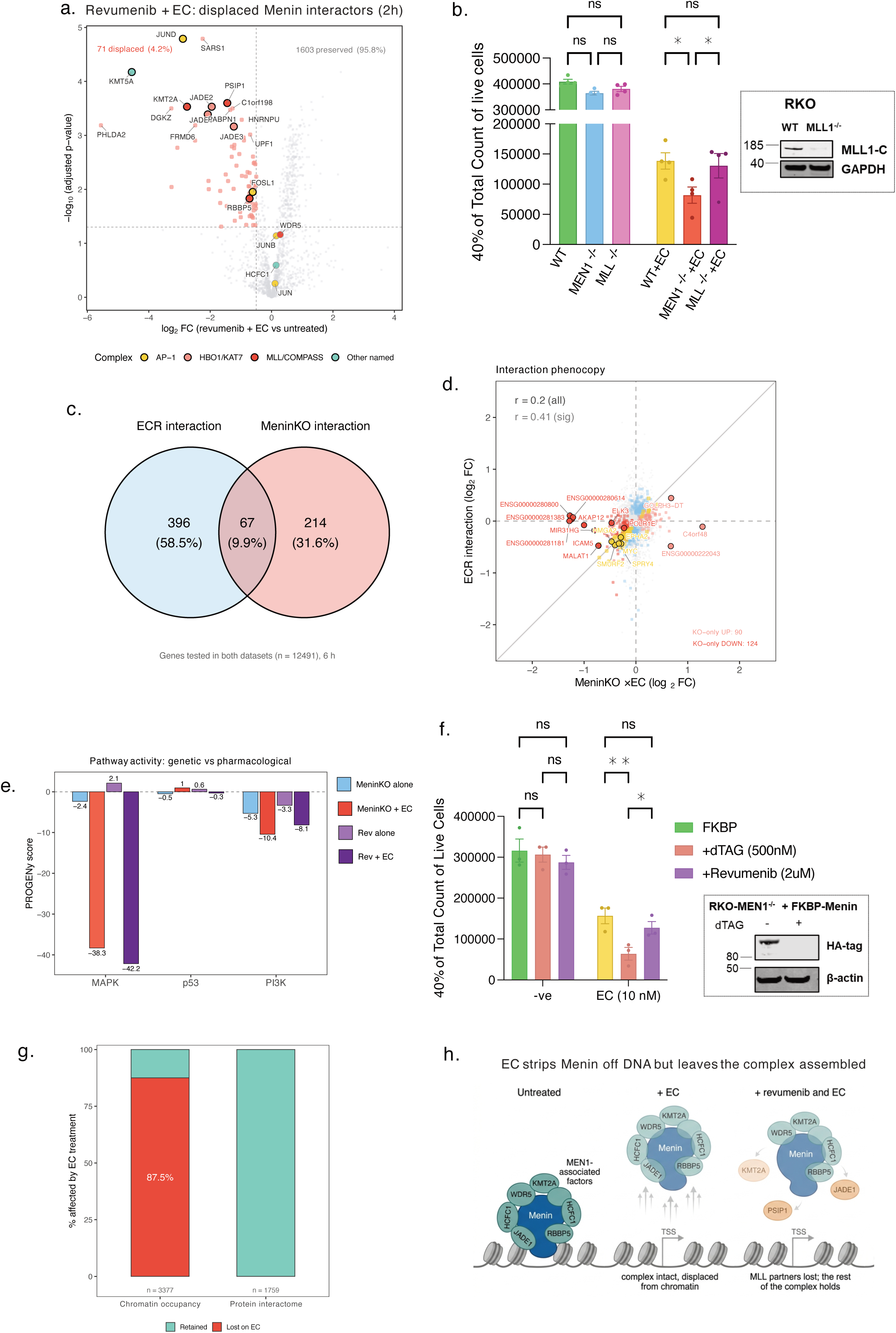
Sensitisation to EC requires loss of Menin rather than inhibition of the Menin/MLL interaction. (a) Differential Menin proximity proteomics following revumenib plus EC treatment relative to untreated cells. Proteins displaced from Menin following treatment are indicated, including KMT2A and associated MLL/COMPASS components. Fill indicates complex membership. Displacement was called at adjusted p < 0.05 and log2FC < -0.5. (b) Long-term proliferation assays comparing wild-type, MEN1-knockout and MLL/KMT2A-knockout RKO cells in untreated and EC (500 nM and 20 µg/mL respectively)-treated conditions. Cell abundance was followed over time to compare the effects of MEN1 and KMT2A loss on sensitivity to EC. Data presented as mean ± SEM of four independent experiments (n=4). Statistical significance was determined using two-way ANOVA. ns = not significant, * p < 0.05. (c) Overlap between genes showing an interaction response to EC following MEN1 knockout and genes showing an interaction response following revumenib treatment. Analysis is restricted to genes tested in both datasets. (d) Comparison of gene-level interaction effects produced by MEN1 knockout and revumenib treatment. Genes significantly altered by either perturbation are indicated. Correlation is reported for all tested genes and separately for genes significant in either contrast. (e) PROGENy pathway activity following MEN1 knockout or revumenib treatment, alone or in combination with EC, showing inferred changes in MAPK, p53 and PI3K pathway activity. Scores are direct contrasts against untreated WT cells at 6h. (f) Viable-cell counts following acute Menin degradation using the FKBP/dTAG (500 nM) system or treatment with revumenib (2 µM), with or without EC (10 nM and 20 µg/mL respectively). Menin degradation, but not inhibition of the Menin/MLL interaction with revumenib, sensitised cells to EC. Data presented as mean ± SEM of three independent experiments (n=3). Statistical significance was determined using two-way ANOVA. ns = not significant, * p < 0.05, **p < 0.01. (g) Comparison of the fraction of Menin-dependent chromatin sites showing reduced occupancy following EC treatment (log2FC < 0) with the fraction of quantified Menin-interacting proteins displaced following treatment (p. adj < 0.05, log2FC < -0.5) (h) Summary model showing the effects of EC on Menin chromatin occupancy and the Menin-associated protein complex.

Genetically, long-term proliferation assays showed that *MEN1*⁻/⁻, *MLL*⁻/⁻ and wild-type cells were indistinguishable in the absence of treatment, but that EC selectively depleted *MEN1*⁻/⁻ cells relative to both wild-type and *MLL*⁻/⁻ cells, while *MLL*⁻/⁻ cells were indistinguishable from wild-type (**Figure 4b**). The MLL arm of Menin function is therefore dispensable for sensitisation to EC.

The transcriptional consequences of the two perturbations also diverged. Comparing the interaction response to EC under *MEN1* knockout with that under revumenib, restricted to the 12,491 genes tested in both datasets, only 67 genes were shared with 9.9% of those responding to either (**Figure 4c**). The transcriptional consequences of the two perturbations also diverged. Among the 12,491 genes tested in both datasets, only 67 genes were differentially expressed in response to EC under both MEN1 knockout and revumenib treatment, representing 9.9% of genes that responded significantly to either perturbation (Figure 4c). Consistent with this limited overlap, the interaction responses showed only modest correlation (r = 0.20 across all tested genes and r = 0.41 among genes significant in either contrast; Figure 4d). Together, these findings indicate that genetic loss of MEN1 and pharmacological inhibition of Menin produce largely distinct transcriptional responses to EC. Genes responding only to Menin loss comprised an induced innate immune module including Toll-like receptor 9 signalling, interleukin-12 production, dendritic cell chemotaxis, together with a repressed actin cytoskeletal and extracellular matrix program. Genes responding only to revumenib were dominated by a strongly repressed ribosome biogenesis signature spanning rRNA processing and ribosomal subunit assembly, which were the most significantly enriched block in the analysis (**Supplementary Figure 3a).**

Pathway attribution separated the two perturbations at baseline. Revumenib alone did not suppress MAPK activity (score +2.1), whereas Menin loss alone did (−2.4). In combination with EC, MAPK was suppressed comparably by both. The axis that modestly distinguished them was PI3K, suppressed more deeply by genetic loss than by revumenib both alone (−5.3 versus -3.3) and in combination with EC (−10.4 versus -8.1; **Figure 4e**), the same axis most affected by Menin loss in **Figure 2a**.

Finally, we tested targeted degradation of Menin. dTAG-mediated degradation of Menin significantly reduced survival under EC relative to both untreated control and revumenib, whereas revumenib produced no significant depletion despite displacing KMT2A and its associated subunits from Menin (**Figure 4a**). Genetic ablation of MLL and pharmacological blockade of the Menin/MLL interface therefore converge on the same negative result, while genetic ablation and targeted degradation of Menin converge on the same positive one. The determinant of sensitisation therefore is the presence of the Menin protein, not the integrity of its MLL binding interface (**Figure 4f**).

Set against one another, two measurements describe complementary effects of EC in which 87% of Menin-dependent chromatin sites lost occupancy, while quantified interactome proteins were not displaced (**Figure 4g**). EC therefore does not diminish Menin per se, it relocates it. The protein may be stripped from DNA with its complex fully assembled, leaving an intact Menin platform that no longer occupies chromatin. Revumenib produces the opposite partial state, dismantling the MLL interface while leaving occupancy untouched, and neither perturbation is lethal. Only degradation removes both, and only degradation phenocopies *MEN1* deletion. The synthetic lethality is not a matter of how much Menin function remains, but of which function remains: EC and revumenib each leave a different half standing, and cells survive on either one (**Figure 4h**).

## Discussion

Here, we identify MEN1 as a factor that determines sensitivity to MAPK pathway inhibition in BRAF-mutant colorectal cancer cells. MEN1 was identified in genome-wide CRISPR screens with both encorafenib plus cetuximab (EC) and ERK inhibition, despite not being a general dependency in CRC cells. MEN1 knockout increased sensitivity to treatment, and this was reversed by re-expression of Menin. This suggests that Menin has an important role in the response to MAPK inhibition rather than simply being required for CRC cell survival.

The screen itself points towards the observed endpoint, phenotypically and transcriptionally. MAPK14, which encodes p38α, provides a stress activated survival signal when canonical ERK signalling is blocked, and MCL1 sequesters BAX and BAK at the mitochondrial membrane. FBXO11, another top sensitiser in both the EC and ERKi screens neddylates p53 and suppresses its transcriptional activity without altering its protein abundance. The prevalence of these hits as sensitisers is consistent with the transcriptional programs observed in MEN1 deficient cells.

ChIP-seq data also suggests that this effect is linked to MAPK-dependent transcription. Menin was bound at a number of MAPK/BRAF-responsive genes, including DUSP5, EGR1, ETV5 and MYC. EC treatment caused a marked global loss of Menin from chromatin. However, the loss of Menin from promoter proximal sites at MAPK/BRAF-responsive genes suggests that MAPK signalling influences how Menin engages with actively regulated genes. This fits with the genetic data, where the effect of MEN1 loss becomes most apparent when MAPK signalling is inhibited.

The proteomic data help separate the loss of Menin from chromatin from disruption of the Menin complex itself. Although EC caused widespread changes in phosphorylation and a substantial reduction in Menin chromatin occupancy, most proteins in the Menin proximity interactome were retained. These included known Menin associated proteins such as KMT2A, WDR5, RBBP5, HCFC1 and PSIP1. EC therefore appears to change Menin chromatin engagement without causing broad disruption of its protein interactions. This may be an important part of how BRAF-mutant cells compensate for MAPK inhibition.

The distinction between MEN1 loss and inhibition of the Menin-MLL interaction was also evident. MLL1 knockout did not reproduce the effect of MEN1 knockout, and revumenib did not sensitise cells to EC to the same extent as MEN1 loss. In contrast, acute degradation of Menin increased sensitivity to EC. The simplest interpretation is that the relevant function of Menin in this setting is not dependent on its interaction with MLL. It also suggests that removing Menin is functionally different from disrupting the interaction targeted by current Menin-MLL inhibitors^20^.

Our data therefore support a model in which Menin helps BRAF-mutant CRC cells adapt to MAPK pathway inhibition. When MAPK signalling is inhibited, Menin is displaced from a substantial fraction of its chromatin sites and the transcriptional state of the cell is altered. Loss of Menin further compromises this response and increases sensitivity to treatment. The fact that Menin degradation, but not pharmacological disruption of the Menin-MLL1 interaction or MLL1 loss, reproduces this phenotype suggests that other Menin dependent interactions are involved and will be important to define.

More broadly, these results support the development of Menin-specific PROTACs as opposed to Menin-MLL inhibitors for BRAF mutant CRC, and describe a distinction that is not specific to Menin. A chromatin-associated scaffold can be removed from DNA while its complex remains assembled and these are separated druggable states.

## Methods

### Cell culture

RKO human colon carcinoma cells and their derivatives were cultured in Dulbecco’s Modified Eagle Medium: Nutrient Mixture F12 (DMEM/F12) supplemented with 5% heat-inactivated fetal bovine serum, 1% GlutaMAX and 1% penicillin-streptomycin (complete DMEM/F12). HEK293T and Lenti-X 293T cells were cultured in the same medium. Cells expressing the TurboID construct were cultured in high-glucose DMEM supplemented with 5% heat-inactivated dialysed FBS, 1% GlutaMAX and 1% penicillin-streptomycin (biotin-free complete DMEM). All cells were maintained at 37 °C in 5% CO₂ and passaged 1:10 using TrypLE. All cells were obtained from ATCC and routinely tested for mycoplasma.

### Generation of knockout cell lines

Knockouts were generated by ribonucleoprotein nucleofection using the SF Cell Line 4D-Nucleofector X Kit (Lonza). A guide RNA complex was assembled by combining 1.2 µL Alt-R S.p. Cas9 nuclease V3 (10 mg/mL), 1 µL gRNA (200 µM) and 3 µL PBS, and incubated for 10 min at room temperature. Cells (250,000) were washed twice in PBS, resuspended in 20 µL nucleofection buffer, combined with 5 µL of the gRNA complex and nucleofected using programme HEK293 on a 4D-Nucleofector Core Unit. Cells were recovered in complete DMEM/F12 for 72 h before expansion. Editing was confirmed by Sanger sequencing with analysis in Synthego ICE, or by immunoblot. sgRNA sequences are listed in Supplementary Table.

### Molecular cloning and retroviral transduction

Coding sequences were synthesised as double-stranded gene fragments (IDT gBlocks) with flanking EcoRI and XhoI overhangs and cloned into an MSCV-based retroviral vector carrying an IRES-linked fluorescent reporter (GFP or BFP). Vector and insert were digested for 2 h at 37 °C, gel-purified, and ligated overnight at 4 °C at vector-to-insert molar ratios of 1:1, 1:3 and 1:7 alongside no-insert and no-ligase controls. Ligations were transformed into chemically competent *E. coli* JM109, plated on ampicillin-containing LB agar, and single colonies expanded for plasmid preparation and verification by Sanger sequencing.

For retroviral production, 14×10^6^ HEK293T cells were seeded per 175 cm^2^ flask and transfected 24 h later with retroviral packaging vectors using polyethylenimine. Medium was replaced after 16 h, and viral supernatant harvested 24 h later and filtered (0.45 µm). Target cells (3×10^6^ per 175 cm^2^ flask) were transduced for 72 h and sorted for the fluorescent reporter on a FACSAria cell sorter. FLAG-tagged Menin was reintroduced into *MEN1*-knockout RKO cells by this route (RKO-*MEN1* KO-FLAG-Menin).

### Genome-wide CRISPR screen

RKO cells were transduced with FUCas9Cherry lentivirus (Addgene #70182) in the presence of 4 µg/mL sequabrene, and mCherry-positive cells sorted to establish RKO-Cas9.

The human Brunello genome-wide knockout library^31^ (Addgene #73178) was packaged in Lenti-X 293T cells using third-generation lentiviral plasmids, and viral supernatant concentrated with Amicon Ultra-15 filters. The library was titrated against RKO-Cas9 cells across a range of viral volumes, with transduction efficiency assessed by survival after 72 h of puromycin selection (1 µg/mL), and the volume achieving 50% transduction was used for the screen.

RKO-Cas9 cells (100×10^6^) were transduced at this titre with 4 µg/mL sequabrene and selected with 1 µg/mL puromycin for 72 h. Surviving cells were expanded and divided into five populations of 50×10^6^ cells each: a reference population (T0), an untreated arm (DMSO, 1:1000), and arms treated with encorafenib plus cetuximab (EC; 500 nM and 10 µg/mL respectively), AZD0364 (250 nM), or 5-fluorouracil (2.5 µM). Cells were cultured for 15 days across three successive rounds of treatment, with 50×10^6^ cells harvested and stored at each passage, maintaining library representation throughout. Surviving cells were collected as the T_end_ population.

Genomic DNA was isolated from 50×10^6^ cells per condition using the Nucleobond CB 100 kit. The sgRNA-containing region was amplified using a P5 forward primer incorporating an 8-nucleotide stagger and a condition-specific barcoded P7 reverse primer, with approximately 16 parallel reactions per condition to preserve representation. Products were pooled and purified with AMPure XP beads.

### CRISPR screen analysis

Reads were trimmed to remove vector sequence 5′ of the protospacer, anchored on the constant U6 sequence immediately preceding the sgRNA (GAAACACCG). Trimmed reads were counted with MAGeCK (v0.5.9.2)^32^ against the Broad GPP Brunello library reference (77,441 sgRNAs), using mageck count with automatic 5′ trim detection.

Comparisons were performed with mageck test using default parameters, taking each treatment arm as treatment and the time-matched untreated arm (T_end_) as control. Counts were median-normalised, sgRNA-level significance assessed under a negative binomial model, and genes ranked by robust rank aggregation, with gene-level log_2_FC taken as the median across sgRNAs targeting each gene. Sensitisers were defined as genes with significant negative selection (FDR < 0.25) and no significant positive selection, and resistors reciprocally.

Library representation was assessed per sample by the proportion of sgRNAs with zero counts and by the Gini index. The unselected T0 population showed minimal skew (Gini 0.080; 420 zero-count sgRNAs), as did the untreated, EC and ERKi arms after 15 days of culture (Gini 0.126-0.153; 1478-2742 zero-count sgRNAs). The 5-fluorouracil arm showed substantially greater skew (Gini 0.356; 6519 zero-count sgRNAs).

### DepMap dependency analysis

CRISPR gene effect (Chronos) scores were retrieved from DepMap release 22Q2 using the depmap Bioconductor package (v3.19) Colorectal lines were selected by lineage annotation, and BRAF V600E lines (RKO, HT29, COLO205, LS411N, MDST8, HT115) identified by mutation status. Genes with a Chronos score below -0.5 were considered dependencies.

### Competitive growth assays

Wild-type and knockout RKO cells were labelled by retroviral transduction with MSCV-IRES-GFP and MSCV-IRES-BFP respectively and sorted for the reporter. Labelled populations were combined at 50,000 cells each in a 12-well plate and treated with EC (500 nM and 20 µg/mL respectively), AZD0364 (250 nM) or DMSO. The proportion of each population was determined by flow cytometry on days 0, 3, 6, 9, 12 and 15; at each timepoint cells were dissociated with TrypLE, an aliquot analysed, and the remainder reseeded with fresh drug. A representative image and corresponding quantification from one representative experiment are shown. Data are presented as mean ± SD of technical replicates. Three independent biological experiments were performed with similar results. Statistical significance was determined using a two-way ANOVA.

### Long-term proliferation assays

Cells were seeded at 30,000 per well in 24-well plates. Lines carrying doxycycline-inducible constructs were pre-treated with 1 µg/mL doxycycline for 24h. Cells were treated with the indicated drugs and passaged every 3 days, with drug replenished at each passage. On day 15 cells were counted on a FACSymphony flow cytometer. Data are presented as mean ± SEM from 3-4 independent biological experiments. Statistical significance was determined using a two-way ANOVA.

### Quantitative real-time PCR

RKO wild-type and *MEN1*-knockout cells (3×10^5^) were treated for 6h with varying concentrations of encorafenib at a fixed cetuximab concentration. RNA was extracted with the RNeasy Mini Kit and quantified by NanoDrop. Complementary DNA was synthesised from 1000 ng total RNA using the High-Capacity cDNA Reverse Transcription Kit (Applied Biosystems) on a Veriti thermal cycler, diluted 1:10, and 3 µL used per qPCR reaction with Power SYBR Green Master Mix on a ViiA7 Real-Time PCR System. Melt curve analysis was performed to confirm amplicon specificity. Relative expression was calculated by the ΔΔCt method as 2^−ΔΔCt^.

### Immunoblotting

Cells were lysed in RIPA buffer supplemented with cOmplete protease inhibitor cocktail and PhosSTOP, incubated on ice for at least 30 min with intermittent mixing, and cleared by centrifugation at 13,000 rpm for 10 min. Protein concentration was determined by Bradford assay. Equal protein amounts were denatured in LDS sample buffer with reducing agent at 95 °C for 5 min, resolved on 4-12% NuPAGE Bis-Tris gels in MOPS SDS running buffer, and dry-transferred to PVDF using an iBlot 2 system (20 V, 7 min). Membranes were blocked in Intercept blocking buffer for 1h, incubated overnight at 4 °C with primary antibodies in 50% Intercept buffer in PBS-Tween, washed, and incubated with fluorophore-conjugated secondary antibodies. Signal was acquired on an Odyssey CLx imaging system. Antibodies are listed in Supplementary Table.

### RNA sequencing

Total RNA was isolated from 10^5^ cells using the RNeasy Mini Kit and assessed on a 4200 TapeStation. Libraries were prepared from 100 ng total RNA using the TruSeq RNA Library Preparation Kit (Illumina), quality-checked, pooled, and sequenced on a NextSeq 2000 with the P2 v3 reagent kit using 66 bp paired-end reads.

### RNAseq processing and analysis

Paired-end reads were aligned to the hg38 (hg38.analysisSet.fa) using Rsubread 2.18.0, with an index generated by buildindex and alignment performed with align in RNA mode, supplying a GENCODE human annotation as an exon junction reference.

Reads were assigned to genes with featureCounts, using the same GENCODE annotation, counting exon features summarised by gene symbol, in paired-end mode and including multi-mapping reads. Counting was performed without strand specification (unstranded libraries).

Counts were filtered with edgeR (v4.2.2)^33^ retaining genes with log-CPM above log2(10/M + 2/L), where L and M are the mean and median library size in millions, in at least as many samples as the smallest experimental group. For the 72 h dataset genes were retained if this threshold was met in at least two samples. Libraries were normalised by trimmed mean of M-values and transformed with voomWithQualityWeights^34^.

Differential expression was assessed with limma (v3.60.6) using a means model (∼0 + condition) with explicit contrasts and Benjamini-Hochberg adjustment. Two classes of contrast are reported. Direct contrasts compare a single condition against untreated wild-type cells. Interaction terms isolate the non-additive component of the combined perturbation and were computed as (MeninKO_EC − MeninKO_untreated) − (WT_EC − WT_untreated) for the knockout dataset and as ECR − EC − Revumenib + Untreated for the drug dataset.

Comparisons between the knockout and drug datasets were restricted to the 12,491 genes tested in both, so that genes excluded from one dataset by expression filtering were not scored as uniquely responsive in the other. RNA-seq data is available (GSE343908).

### Pathway activity inference

Pathway activities were inferred with PROGENy via decoupleR (v2.10.0) using a univariate linear model with the top 500 targets per pathway and a minimum set size of 5, taking moderated t-statistics from the relevant limma contrast as input. PROGENy scores pathway activity from the coordinated expression of experimentally derived target genes rather than from the expression of pathway components, so a pathway can register altered activity without any change in the abundance of its own transcripts.

Where activities from independent experiments were compared, input t-statistics were z-scaled within each dataset before inference; such comparisons are interpreted on direction and rank rather than absolute magnitude. Per-gene contributions to a pathway score were calculated as the product of the moderated t-statistic and the PROGENy regulon weight.

### Gene set enrichment

GO Biological Process enrichment was performed with clusterProfiler [version], applying Benjamini-Hochberg adjustment and a set size range of 5-500, analysing genes responding uniquely to each perturbation separately by direction against the 12,491 genes tested in both RNA-seq datasets as background.

### ChIP sequencing

RKO wild-type and *MEN1*⁻/⁻ cells were treated with encorafenib (500 nM) and cetuximab (20 µg/mL) or vehicle for 6h. ChIP-seq was performed as previously described (Vervoort et al., 2021). Cells (10 × 10^6^ per immunoprecipitation) were washed once in cold PBS and cross-linked at room temperature for 10 min with one-tenth volume of freshly prepared formaldehyde solution (11% formaldehyde, 0.5 mM EGTA, 1 mM EDTA, 100 mM NaCl, 50 mM HEPES-KOH pH 7.5) in PBS at room temperature. Cross-linking was quenched by addition of one-twentieth volume of 2.5 M glycine followed by a further 5 min at room temperature. Cells were washed in cold PBS, scraped and pelleted. From scraping onwards all buffers were supplemented with cOmplete EDTA-free protease inhibitor cocktail and PhosSTOP (Roche).

Nuclei were isolated by three successive 10 min incubations on ice in cold nuclear extraction buffer (20 mM Tris-HCl pH 8, 10 mM NaCl, 0.5% IGEPAL CA-630, 2 mM EDTA), resuspended in sonication buffer (20 mM Tris-HCl pH 7.5, 150 mM NaCl, 2 mM EDTA, 0.3% SDS, 1% IGEPAL CA-630) and sonicated for 8 min on a Covaris ME220 (peak power 75, duty factor 15%, 1,000 cycles per burst, average power 11.3). Fragmented chromatin was cleared by centrifugation at maximum speed for 20 min at 4 °C and the supernatant diluted 1:1 in dilution buffer (20 mM Tris-HCl pH 8, 150 mM NaCl, 2 mM EDTA, 1% Triton X-100). Chromatin was quantified using the Qubit dsDNA BR assay (Invitrogen) and NIH-3T3 (mouse) chromatin was spiked into each sample at 3% of total chromatin mass.

For each immunoprecipitation, diluted chromatin was rotated overnight at 4 °C with Pierce Protein A/G magnetic beads (Thermo Scientific) and 1 µg anti-Menin antibody(Cell Signaling Technologies #19893). Beads were washed once each in ChIP buffer (20 mM Tris-HCl pH 8, 150 mM NaCl, 2 mM EDTA, 0.15% SDS, 1% Triton X-100), ChIP wash buffer 1 (20 mM Tris-HCl pH 8, 500 mM NaCl, 2 mM EDTA, 0.1% SDS, 1% Triton X-100) and ChIP wash buffer 2 (20 mM Tris-HCl pH 8, 250 mM LiCl, 2 mM EDTA, 0.5% deoxycholate, 0.5% IGEPAL CA-630), then twice in TE buffer (10 mM Tris-HCl pH 7.5, 1 mM EDTA). Washed beads were incubated with shaking at 55 °C for 1 h in reverse cross-linking buffer (200 mM NaCl, 100 mM NaHCO_3_, 1% SDS, 300 µg/mL proteinase K), and the supernatant incubated overnight at 65 °C. DNA was purified using the ChIP DNA Clean and Concentrator Kit (Zymo Research D5205) according to the manufacturer’s instructions. Sequencing libraries were prepared with the NEBNext Ultra II DNA Library Prep Kit (NEB E7645) and sequenced on Element AVITI24 to a depth of 20 million reads per sample. One replicate was performed per condition. ChIP-seq data is available (GSE343967).

### ChIP-seq processing and analysis

Paired-end reads were aligned with bowtie2 2.5.3^35^ using --local --sensitive-local --no-mixed -- no-discordant --phred33 -I 10 -X 700, which restricts to concordantly aligned pairs with inserts sizes between 10-700bp. Reads were aligned in parallel to the hg38 human genome and the mm10 mouse genome for the spike-in. Alignments were coordinate-sorted with samtools (v1.23.2) and duplicates removed with Picard MarkDuplicates 2.27.5 (REMOVE_DUPLICATES=true).

Peaks were called with MACS2 (v2.2.9.1)^36^ using callpeak -f BAMPE -g hs -q 0.01 --keep-dup all, each immunoprecipitation against its own matched input. Peaks were restricted to primary contigs (chr1-22, X, Y). Coverage tracks were generated with deepTools^37^ bamCoverage 3.5.5 at 10 bp bin resolution and normalised to counts per million (−-normalizeUsing CPM). Reference-adjusted (ChIP-Rx) tracks, scaled by the reciprocal of each sample’s mm10 spike-in read count in millions, were also generated but were not used for the analyses reported here. Signal was quantified with deepTools multiBigwigSummary from the CPM tracks, at the Menin-dependent sites (BED-file mode) and genome-wide in 10 kb bins (bins mode). The genome wied bins provide a background against which change at menin dependent sites can be assessed and were restricted to bins with signal in at least one condition. Sites were annotated to the nearest protein-coding transcription start site with bedtools (v2.31.1) closest

Peaks called in *MEN1*⁻/⁻ cells were used to define antibody background. A Menin-dependent site was defined as a peak present in wild-type cells and absent in *MEN1*⁻/⁻ cells (bedtools intersect -v). Of 4,919 peaks called in wild-type cells on primary contigs, 1,542 overlapped a peak called in *MEN1*⁻/⁻ cells and were excluded; the remaining 3,377 wild-type-specific peaks were used for all subsequent analysis (any WT peak sharing a single base with a KO peak was discarded).

Treatment responses were expressed as log2 ((treated + 0.05)/(untreated + 0.05)); the equivalent ratio in *MEN1*⁻/⁻ cells was calculated in parallel as a control for treatment effects independent of Menin; so a site losing signal in KO where no menin is expected to be present cannot have menin displaced. Sites equidistant from two annotated start sites were assigned to a single gene. Sites within 2 kb of a transcription start site were classified as promoter-proximal, yielding 1,607 Menin-bound promoter genes; where a gene carried several promoter-proximal sites, the site with the largest absolute treatment response was used. Genome-wide background was computed in 10 kb bins with signal in at least one condition, and distributions compared by two-sided Kolmogorov-Smirnov test.

### Proximity labelling

RKO *MEN1*-knockout cells carrying a doxycycline-inducible TurboID-Menin construct (RKO-*MEN1* KO-TurboID-Menin) were seeded at 7.5 × 10^6^ cells in biotin-free complete DMEM and cultured with or without 1 µg/mL doxycycline for 24 h. Cells were then treated with the indicated drugs for 2 h, with 100 µM biotin added for the final 40 min of treatment. Because the construct is expressed in a *MEN1*-knockout background, the TurboID-Menin fusion replaces rather than supplements endogenous Menin. Cells were washed five times in ice-cold PBS and lysed in 1 mL RIPA buffer supplemented with 0.1% SDS, benzonase, protease inhibitor and PhosSTOP, with end-over-end rotation for 1 h at 4 °C. Lysates were cleared at 20,000 × g for 15 min and protein quantified using the Qubit protein assay. Biotinylated proteins were enriched on Pierce high-capacity streptavidin agarose beads (10 µL slurry per sample) for 60 min at 4 °C with rotation, and beads washed three times in PBS containing 0.5% SDS. Proteins were reduced in 100 mM DTT for 20 min, washed into 6 M urea/100 mM Tris-HCl pH 8.5, alkylated with 50 mM iodoacetamide for 20 min in the dark, then washed ten times in urea buffer, four times in PBS and three times in water. On-bead digestion was performed overnight at 37 °C in 50 mM ammonium bicarbonate with 2 µg Trypsin Gold. Peptides were recovered by centrifugation, beads washed once, fractions pooled, dried and reconstituted in loading buffer. Peptides were separated on a 15 cm C18 fused-silica column with integrated emitter tip (IonOpticks; 75 µm ID, 1.6 µm beads) using a Vanquish Neo UHPLC coupled to an Orbitrap Eclipse Tribrid mass spectrometer. Peptides were loaded at 600 nL/min in buffer A (0.1% formic acid) and eluted at 400 nL/min over a 30 min linear gradient of 2-34% buffer B (80% acetonitrile, 0.1% formic acid). Data were acquired in data-independent mode: MS1 in the Orbitrap at 120,000 resolution (normalised AGC 300%, RF lens 40%, scan range 375-1500 m/z), MS2 in the Orbitrap at 30,000 resolution (first mass 120 m/z, normalised AGC 100%, maximum injection time 54 ms).

### Proximity labelling analysis

Proteins were included in the Menin interactome only if significantly enriched over both controls (cells with biotin without doxycycline induction, and cells induced with doxycycline but no biotin added) at Benjamini-Hochberg adjusted p < 0.01 and log2FC > 2 in each comparison. This yielded 1,759 proteins. Recovery of previously reported interactors was assessed against the BioGRID *MEN1* record (BIOGRID-GENE-110384, release 5.0.258) restricted to human physical interactions. Displacement under treatment was called at adjusted p < 0.05 and log2FC < -0.5 among interactome members quantified in the relevant comparison; percentages are expressed relative to the number of proteins quantified in that comparison.

### Phosphoproteomics

Cells were lysed in 200 µL ice-cold SDC buffer (4% w/v sodium deoxycholate, 100 mM Tris-HCl pH 8.5), heated to 95 °C for 5 min, sonicated for 15 s with a tip probe and cleared at 13,000 rpm for 10 min at 4 °C. Protein was quantified by BCA assay. Protein (500 µg per sample) was reduced and alkylated in 100 mM TCEP/400 mM chloroacetamide for 5 min at 45 °C, then digested overnight at 37 °C with Lys-C and trypsin at an enzyme-to-protein ratio of 1:100.

Phosphopeptides were enriched on titanium dioxide beads (Titansphere Phos-TiO, 6 mg per sample) following addition of isopropanol and enrichment buffer, with an aliquot retained for total proteome analysis. Beads were washed five times in 5% trifluoroacetic acid/8 mM KH₂PO₄, transferred to in-house C8 StageTips, and phosphopeptides eluted in 40% acetonitrile with ammonium hydroxide. Eluates were dried, resuspended in 1% trifluoroacetic acid/99% isopropanol, desalted on SDB-RPS StageTips, eluted in 60% acetonitrile with ammonium hydroxide, dried and reconstituted in loading buffer.

Peptides were separated on a 15 cm C18 fused-silica column with integrated emitter tip (IonOpticks; 75 µm ID, 1.6 µm beads) using a Vanquish Neo UHPLC coupled to an Orbitrap Astral mass spectrometer, over a 23 min linear gradient of 2-34% buffer B (80% acetonitrile, 0.1% formic acid). Data were acquired in data-independent mode: MS1 at 240,000 resolution across 380-980 m/z with a normalised AGC target of 500%; MS2 with 4 m/z isolation windows and higher-energy collisional dissociation at a normalised collision energy of 25%, detecting fragments across 145-1450 m/z with a maximum injection time of 6 ms and normalised AGC target of 800%.

Raw data were processed in Spectronaut (version 19.9) using the directDIA workflow with default settings, specifying cysteine carbamidomethylation as a fixed modification and phosphorylation of serine, threonine and tyrosine, methionine oxidation and N-terminal acetylation as variable modifications. Searches were performed against the *Homo sapiens*reference proteome with phosphosite localisation enabled. Precursor and protein Q-value and posterior error probability cutoffs were set to 0.01. Spectronaut output was collapsed to phosphosites with a localisation threshold of 0.75 using the peptide collapse plug-in. Phosphosite intensities were log₂-transformed and filtered to retain sites quantified in all four biological replicates, with missing values imputed from a normal distribution (width 0.3, downshift 1.8) in Perseus (version 1.6.2.3).

### Phosphoproteomics analysis

Phosphosites with an assigned single residue and a reported fold change and q-value for the EC versus untreated comparison were retained, and significance called at q < 0.05. Domain diagrams were drawn against UniProt sequence annotations (accessions P00533, P15056, P36507, P28482, P27361, Q15418, P01106, P05412, P29317). Comparisons with the Menin interactome were restricted to proteins with at least one measured phosphosite and tested by Fisher’s exact test against that background. Because phosphosites rather than proteins are the unit of measurement, a protein may contribute to both the increased and decreased sets.

### Data visualisation and statistics

Expression heatmaps display log2 CPM centred on the mean of each gene across all samples, without scaling to unit variance, so that colour is expressed in log2 units and is comparable between rows. Statistical tests for individual experiments are specified in the corresponding figure legends.

## Acknowledgements

We thank the WEHI genomics facility and the WEHI proteomics facility. C.J.K. is supported by an NHMRC Investigator grant EL2 (2034017), an NHMRC Ideas grant (2029625), and a project grant from Tour de Cure. S.J.V. was partially/fully funded by the Snow Medical Research Foundation through the support of the Snow Fellowship program and a CSL Centenary fellowship.

## Author Contributions

JM, CJK and SJV designed and supervised the study. AS performed the majority of experiments and AAH performed experiments and the majority of data analysis and figure preparation.

## Competing Interests

The authors declare no competing intertests.

---

**Supplementary Figure 1.**
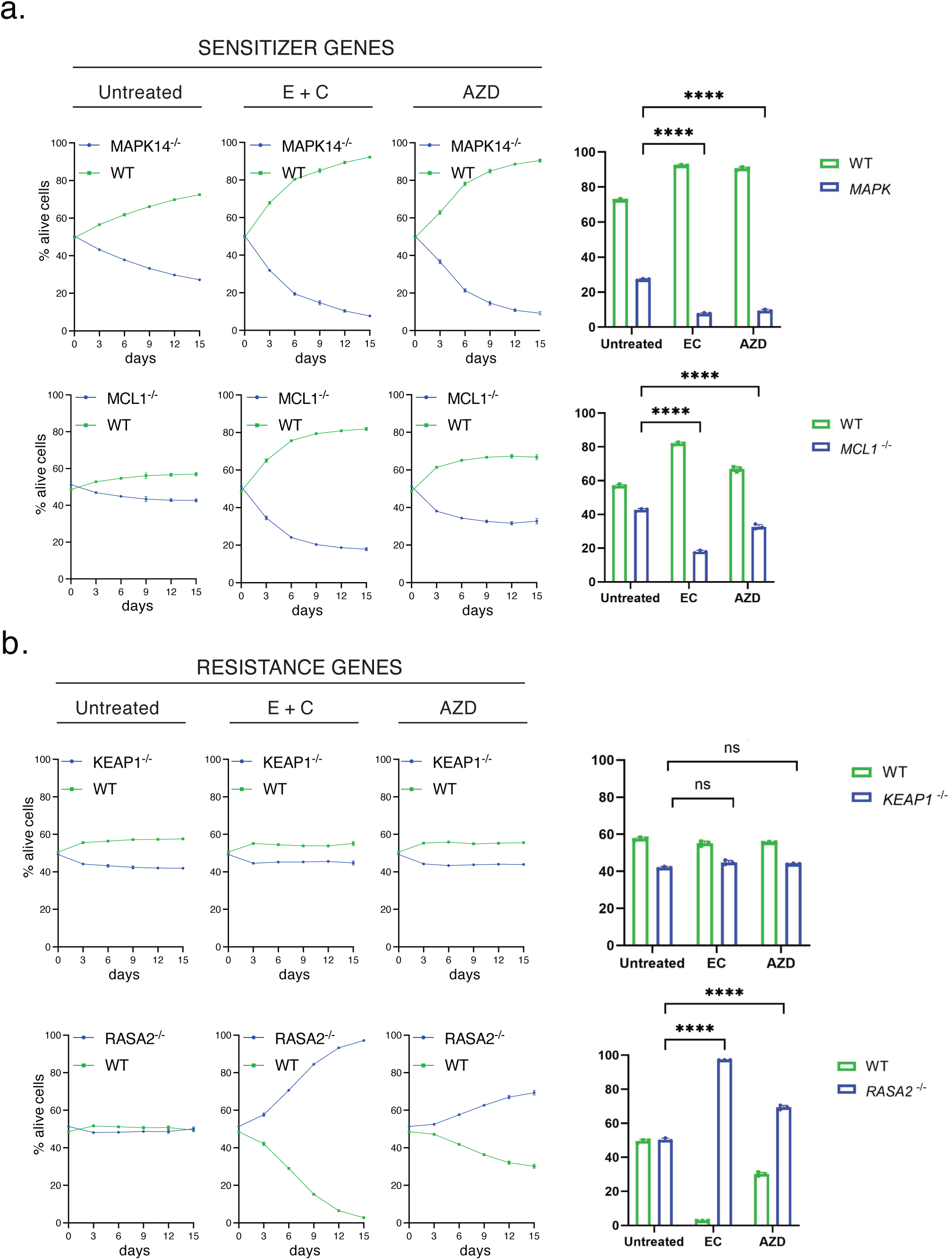
Validation of sensitiser and resistance genes identified by CRISPR screening. (a) Competition assays comparing wild-type cells with MAPK14 or MCL1-knockout cells under untreated, EC(500 nM and 20 µg/mL respectively)- and ERKi (250 nM)-treated conditions. The proportion of viable cells of each genotype was followed over time, with endpoint quantification shown at right. Representative images from one of the three independent experiments (n=3) are shown. Data presented as mean ± SD. Statistical significance was determined using two-way ANOVA. ****p < 0.0001. (b) Competition assays comparing wild-type cells with KEAP1 or RASA2-knockout cells under untreated, EC (500 nM and 20 µg/mL respectively) and ERKi (250 nM)-treated conditions. The proportion of viable cells of each genotype was followed over time, with endpoint quantification shown at right. Representative images from one of the three independent experiments (n=3) are shown. Data presented as mean ± SD. Statistical significance was determined using two-way ANOVA. ns = not significant, ****p < 0.0001.

**Supplementary Figure 2.**
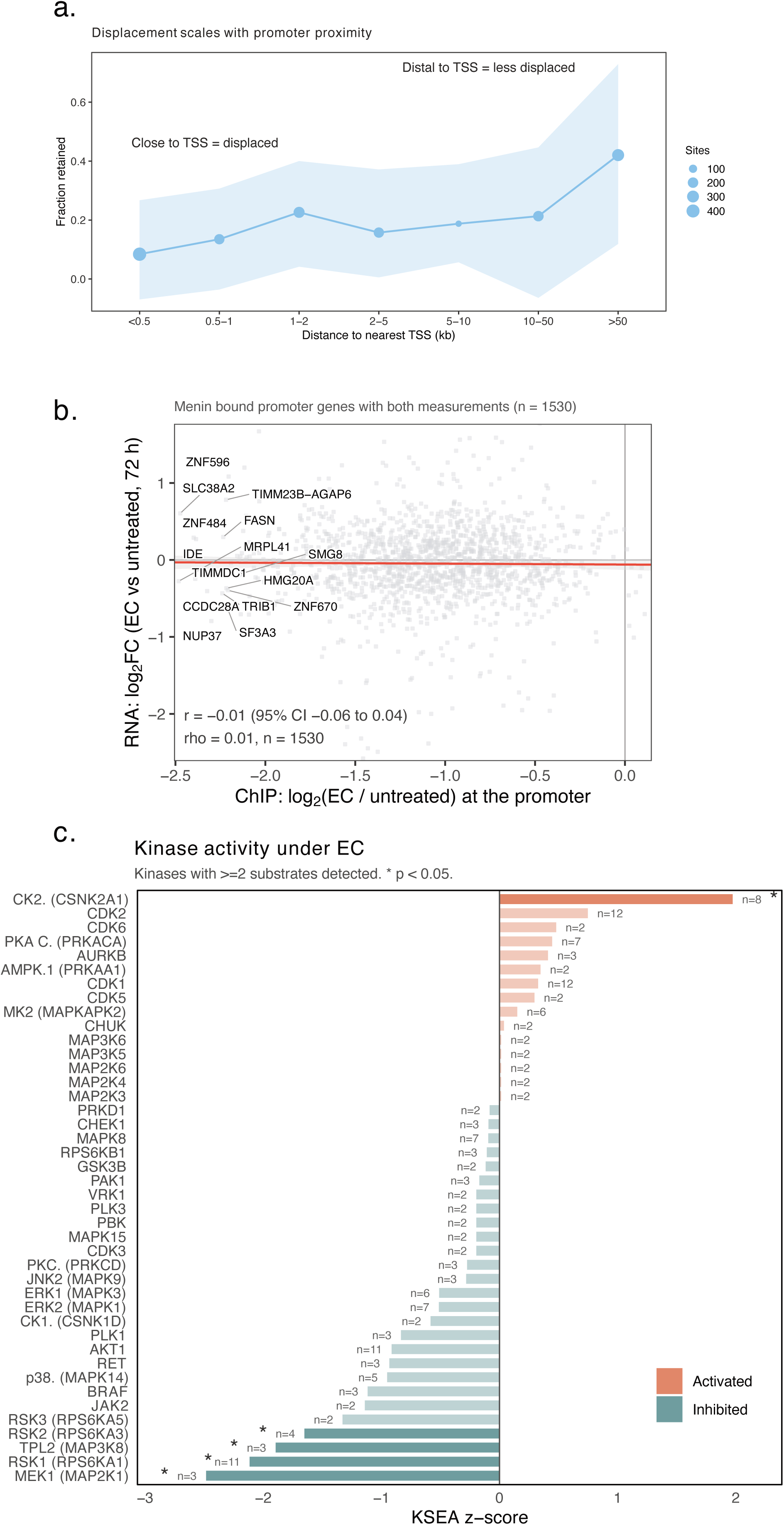
Additional analysis of Menin chromatin occupancy, signalling and protein interactions following EC treatment. (a) Fraction of Menin chromatin occupancy retained following EC treatment binned by distance to the nearest transcription start site; point size, sites per bin; sahded band, 95% CI. (b) Relationship between the EC-induced change in Menin occupancy (log2FC) at Menin-bound promoters and the corresponding transcriptional response (EC vs untreated, 72h). Red line, linear fit with 95% confidence band; the fifteen most displaced genes are labelled. (c) Kinase-substrate enrichment analysis of the phosphoproteomic response to EC. Kinases represented by at least two quantified substrates are shown. For each kinase, z-score compares the mean log2FC of its annotated substrate against all quantified phosphosites,

**Supplementary Figure 3.**
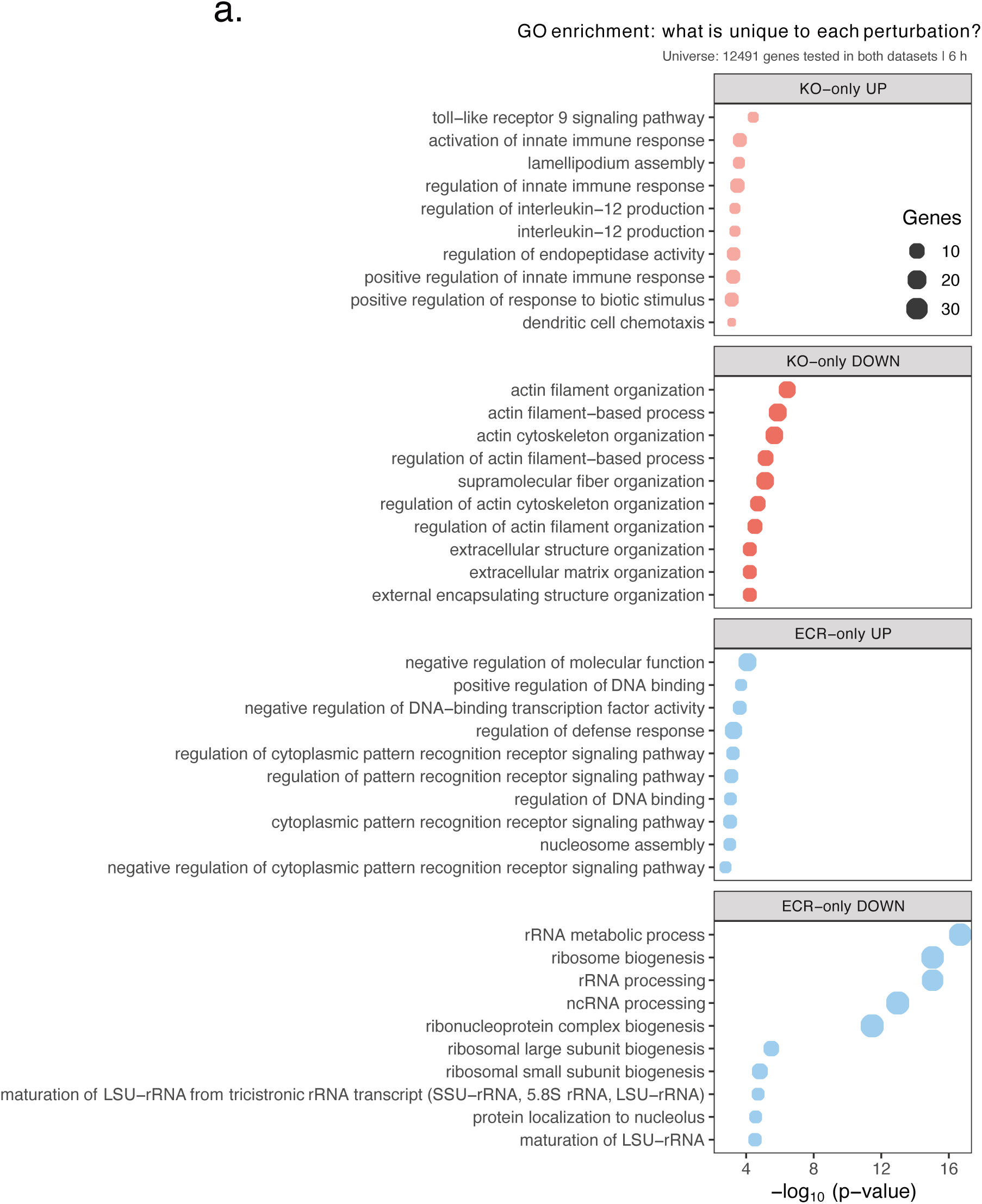
Gene ontology analysis of transcriptional responses unique to MEN1 loss or revumenib treatment. (a) Gene Ontology Biological Process enrichment analysis of genes uniquely increased or decreased following MEN1 knockout or revumenib treatment in the context of EC. Upregulated and downregulated genes were analysed separately for each perturbation. Dot size indicates the number of genes associated with each term and the x-axis shows -log10(P value). The background comprised the 12,491 genes tested in both RNA-seq datasets.

## Notes

### Competing Interest Statement

The authors have declared no competing interest.

## References

1. Van Cutsem, E., Kohne, C.H., Lang, I., Folprecht, G., Nowacki, M.P., Cascinu, S., Shchepotin, I., Maurel, J., Cunningham, D., Tejpar, S., et al. (2011). Cetuximab plus irinotecan, fluorouracil, and leucovorin as first-line treatment for metastatic colorectal cancer: updated analysis of overall survival according to tumor KRAS and BRAF mutation status. J Clin Oncol 29, 2011–2019. 10.1200/JCO.2010.33.5091.

2. Prahallad, A., Sun, C., Huang, S., Di Nicolantonio, F., Salazar, R., Zecchin, D., Beijersbergen, R.L., Bardelli, A., and Bernards, R. (2012). Unresponsiveness of colon cancer to BRAF(V600E) inhibition through feedback activation of EGFR. Nature 483, 100–103. 10.1038/nature10868.

3. Corcoran, R.B., Ebi, H., Turke, A.B., Coffee, E.M., Nishino, M., Cogdill, A.P., Brown, R.D., Della Pelle, P., Dias-Santagata, D., Hung, K.E., et al. (2012). EGFR-mediated re-activation of MAPK signaling contributes to insensitivity of BRAF mutant colorectal cancers to RAF inhibition with vemurafenib. Cancer Discov 2, 227–235. 10.1158/2159-8290.CD-11-0341.

4. Corcoran, R.B., Andre, T., Atreya, C.E., Schellens, J.H.M., Yoshino, T., Bendell, J.C., Hollebecque, A., McRee, A.J., Siena, S., Middleton, G., et al. (2018). Combined BRAF, EGFR, and MEK Inhibition in Patients with BRAF(V600E)-Mutant Colorectal Cancer. Cancer Discov 8, 428–443. 10.1158/2159-8290.CD-17-1226.

5. van Geel, R., Tabernero, J., Elez, E., Bendell, J.C., Spreafico, A., Schuler, M., Yoshino, T., Delord, J.P., Yamada, Y., Lolkema, M.P., et al. (2017). A Phase Ib Dose-Escalation Study of Encorafenib and Cetuximab with or without Alpelisib in Metastatic BRAF-Mutant Colorectal Cancer. Cancer Discov 7, 610–619. 10.1158/2159-8290.CD-16-0795.

6. Kopetz, S., Grothey, A., and Tabernero, J. (2020). Encorafenib, Binimetinib, and Cetuximab in BRAF V600E-Mutated Colorectal Cancer. Reply. N Engl J Med 382, 877–878. 10.1056/NEJMc1915676.

7. Tabernero, J., Grothey, A., Van Cutsem, E., Yaeger, R., Wasan, H., Yoshino, T., Desai, J., Ciardiello, F., Loupakis, F., Hong, Y.S., et al. (2021). Encorafenib Plus Cetuximab as a New Standard of Care for Previously Treated BRAF V600E-Mutant Metastatic Colorectal Cancer: Updated Survival Results and Subgroup Analyses from the BEACON Study. J Clin Oncol 39, 273–284. 10.1200/JCO.20.02088.

8. Kopetz, S., Grothey, A., Yaeger, R., Van Cutsem, E., Desai, J., Yoshino, T., Wasan, H., Ciardiello, F., Loupakis, F., Hong, Y.S., et al. (2019). Encorafenib, Binimetinib, and Cetuximab in BRAF V600E-Mutated Colorectal Cancer. N Engl J Med 381, 1632–1643. 10.1056/NEJMoa1908075.

9. Bock, C., Datlinger, P., Chardon, F., Coelho, M.A., Dong, M.B., Lawson, K.A., Lu, T., Maroc, L., Norman, T.M., Song, B., et al. (2022). High-content CRISPR screening. Nat Rev Methods Primers 2. 10.1038/s43586-022-00098-7.

10. Yang, H., Higgins, B., Kolinsky, K., Packman, K., Bradley, W.D., Lee, R.J., Schostack, K., Simcox, M.E., Kopetz, S., Heimbrook, D., et al. (2012). Antitumor activity of BRAF inhibitor vemurafenib in preclinical models of BRAF-mutant colorectal cancer. Cancer Res 72, 779–789. 10.1158/0008-5472.CAN-11-2941.

11. Chandrasekharappa, S.C., Guru, S.C., Manickam, P., Olufemi, S.E., Collins, F.S., Emmert-Buck, M.R., Debelenko, L.V., Zhuang, Z., Lubensky, I.A., Liotta, L.A., et al. (1997). Positional cloning of the gene for multiple endocrine neoplasia-type 1. Science 276, 404–407. 10.1126/science.276.5311.404.

12. Lemos, M.C., and Thakker, R.V. (2008). Multiple endocrine neoplasia type 1 (MEN1): analysis of 1336 mutations reported in the first decade following identification of the gene. Hum Mutat 29, 22–32. 10.1002/humu.20605.

13. Milne, T.A., Hughes, C.M., Lloyd, R., Yang, Z., Rozenblatt-Rosen, O., Dou, Y., Schnepp, R.W., Krankel, C., Livolsi, V.A., Gibbs, D., et al. (2005). Menin and MLL cooperatively regulate expression of cyclin-dependent kinase inhibitors. Proc Natl Acad Sci U S A 102, 749–754. 10.1073/pnas.0408836102.

14. Agarwal, S.K., Guru, S.C., Heppner, C., Erdos, M.R., Collins, R.M., Park, S.Y., Saggar, S., Chandrasekharappa, S.C., Collins, F.S., Spiegel, A.M., et al. (1999). Menin interacts with the AP1 transcription factor JunD and represses JunD-activated transcription. Cell 96, 143–152. 10.1016/s0092-8674(00)80967-8.

15. Yokoyama, A., Somervaille, T.C., Smith, K.S., Rozenblatt-Rosen, O., Meyerson, M., and Cleary, M.L. (2005). The menin tumor suppressor protein is an essential oncogenic cofactor for MLL-associated leukemogenesis. Cell 123, 207–218. 10.1016/j.cell.2005.09.025.

16. Yokoyama, A., and Cleary, M.L. (2008). Menin critically links MLL proteins with LEDGF on cancer-associated target genes. Cancer Cell 14, 36–46. 10.1016/j.ccr.2008.05.003.

17. Huang, J., Gurung, B., Wan, B., Matkar, S., Veniaminova, N.A., Wan, K., Merchant, J.L., Hua, X., and Lei, M. (2012). The same pocket in menin binds both MLL and JUND but has opposite effects on transcription. Nature 482, 542–546. 10.1038/nature10806.

18. Katona, B.W., Glynn, R.A., Paulosky, K.E., Feng, Z., Davis, C.I., Ma, J., Berry, C.T., Szigety, K.M., Matkar, S., Liu, Y., et al. (2019). Combined Menin and EGFR Inhibitors Synergize to Suppress Colorectal Cancer via EGFR-Independent and Calcium-Mediated Repression of SKP2 Transcription. Cancer Res 79, 2195–2207. 10.1158/0008-5472.CAN-18-2133.

19. Katona, B.W., Hojnacki, T., Glynn, R.A., Paulosky, K.E., Szigety, K.M., Cao, Y., Zhang, X., Feng, Z., He, X., Ma, J., and Hua, X. (2020). Menin-mediated Repression of Glycolysis in Combination with Autophagy Protects Colon Cancer Against Small-molecule EGFR Inhibitors. Mol Cancer Ther 19, 2319–2329. 10.1158/1535-7163.MCT-20-0101.

20. Krivtsov, A.V., Evans, K., Gadrey, J.Y., Eschle, B.K., Hatton, C., Uckelmann, H.J., Ross, K.N., Perner, F., Olsen, S.N., Pritchard, T., et al. (2019). A Menin-MLL Inhibitor Induces Specific Chromatin Changes and Eradicates Disease in Models of MLL-Rearranged Leukemia. Cancer Cell 36, 660–673 e611. 10.1016/j.ccell.2019.11.001.

21. Issa, G.C., Aldoss, I., Thirman, M.J., DiPersio, J., Arellano, M., Blachly, J.S., Mannis, G.N., Perl, A., Dickens, D.S., McMahon, C.M., et al. (2025). Menin Inhibition With Revumenib for KMT2A-Rearranged Relapsed or Refractory Acute Leukemia (AUGMENT-101). J Clin Oncol 43, 75–84. 10.1200/JCO.24.00826.

22. Arellano, M.L., Thirman, M.J., DiPersio, J.F., Heiblig, M., Stein, E.M., Schuh, A.C., Zucenka, A., de Botton, S., Grove, C.S., Mannis, G.N., et al. (2025). Menin inhibition with revumenib for NPM1-mutated relapsed or refractory acute myeloid leukemia: the AUGMENT-101 study. Blood 146, 1065–1077. 10.1182/blood.2025028357.

23. Wagner, E.F., and Nebreda, A.R. (2009). Signal integration by JNK and p38 MAPK pathways in cancer development. Nat Rev Cancer 9, 537–549. 10.1038/nrc2694.

24. Czabotar, P.E., Lessene, G., Strasser, A., and Adams, J.M. (2014). Control of apoptosis by the BCL-2 protein family: implications for physiology and therapy. Nat Rev Mol Cell Biol 15, 49–63. 10.1038/nrm3722.

25. Milella, M., Falcone, I., Conciatori, F., Cesta Incani, U., Del Curatolo, A., Inzerilli, N., Nuzzo, C.M., Vaccaro, V., Vari, S., Cognetti, F., and Ciuffreda, L. (2015). PTEN: Multiple Functions in Human Malignant Tumors. Front Oncol 5, 24. 10.3389/fonc.2015.00024.

26. Arafeh, R., Qutob, N., Emmanuel, R., Keren-Paz, A., Madore, J., Elkahloun, A., Wilmott, J.S., Gartner, J.J., Di Pizio, A., Winograd-Katz, S., et al. (2015). Recurrent inactivating RASA2 mutations in melanoma. Nat Genet 47, 1408–1410. 10.1038/ng.3427.

27. Rojo de la Vega, M., Chapman, E., and Zhang, D.D. (2018). NRF2 and the Hallmarks of Cancer. Cancer Cell 34, 21–43. 10.1016/j.ccell.2018.03.022.

28. Tsherniak, A., Vazquez, F., Montgomery, P.G., Weir, B.A., Kryukov, G., Cowley, G.S., Gill, S., Harrington, W.F., Pantel, S., Krill-Burger, J.M., et al. (2017). Defining a Cancer Dependency Map. Cell 170, 564–576 e516. 10.1016/j.cell.2017.06.010.

29. Schubert, M., Klinger, B., Klunemann, M., Sieber, A., Uhlitz, F., Sauer, S., Garnett, M.J., Bluthgen, N., and Saez-Rodriguez, J. (2018). Perturbation-response genes reveal signaling footprints in cancer gene expression. Nat Commun 9, 20. 10.1038/s41467-017-02391-6.

30. Branon, T.C., Bosch, J.A., Sanchez, A.D., Udeshi, N.D., Svinkina, T., Carr, S.A., Feldman, J.L., Perrimon, N., and Ting, A.Y. (2018). Efficient proximity labeling in living cells and organisms with TurboID. Nat Biotechnol 36, 880–887. 10.1038/nbt.4201.

31. Doench, J.G., Fusi, N., Sullender, M., Hegde, M., Vaimberg, E.W., Donovan, K.F., Smith, I., Tothova, Z., Wilen, C., Orchard, R., et al. (2016). Optimized sgRNA design to maximize activity and minimize off-target effects of CRISPR-Cas9. Nat Biotechnol 34, 184–191. 10.1038/nbt.3437.

32. Li, W., Xu, H., Xiao, T., Cong, L., Love, M.I., Zhang, F., Irizarry, R.A., Liu, J.S., Brown, M., and Liu, X.S. (2014). MAGeCK enables robust identification of essential genes from genome-scale CRISPR/Cas9 knockout screens. Genome Biol 15, 554. 10.1186/s13059-014-0554-4.

33. Robinson, M.D., McCarthy, D.J., and Smyth, G.K. (2010). edgeR: a Bioconductor package for differential expression analysis of digital gene expression data. Bioinformatics 26, 139–140. 10.1093/bioinformatics/btp616.

34. Ritchie, M.E., Phipson, B., Wu, D., Hu, Y., Law, C.W., Shi, W., and Smyth, G.K. (2015). limma powers differential expression analyses for RNA-sequencing and microarray studies. Nucleic Acids Res 43, e47. 10.1093/nar/gkv007.

35. Langmead, B., and Salzberg, S.L. (2012). Fast gapped-read alignment with Bowtie 2. Nat Methods 9, 357–359. 10.1038/nmeth.1923.

36. Zhang, Y., Liu, T., Meyer, C.A., Eeckhoute, J., Johnson, D.S., Bernstein, B.E., Nusbaum, C., Myers, R.M., Brown, M., Li, W., and Liu, X.S. (2008). Model-based analysis of ChIP-Seq (MACS). Genome Biol 9, R137. 10.1186/gb-2008-9-9-r137.

37. Ramirez, F., Ryan, D.P., Gruning, B., Bhardwaj, V., Kilpert, F., Richter, A.S., Heyne, S., Dundar, F., and Manke, T. (2016). deepTools2: a next generation web server for deep-sequencing data analysis. Nucleic Acids Res 44, W160–165. 10.1093/nar/gkw257.

